# Calcineurin B-mediated Ca^2+^ sensing translates stress signal intensity into the assembly of phase-separated condensates at PERK complexes

**DOI:** 10.64898/2026.08.03.742535

**Authors:** Sebastian M. Bairo, Macarena Fernandez, Gonzalo Quasollo, Andrea Pellegrini, Juan C. de Batista, Santiago Asis, Mauricio Martin, Deborah Holstein, James D. Lechleiter, Gabriela E. Gomez, Mariano Bisbal, Mariana Bollo

## Abstract

Endoplasmic reticulum (ER) stress activates protein kinase RNA-like ER kinase (PERK), which initially promotes adaptive responses but remains the only active UPR branch during prolonged stress, mediating both early cytoprotective and chronic pro-apoptotic signaling. Recently, we identified translocon-generated Ca^2+^ microdomains that promote PERK phosphorylation during early UPR, revealing a mechanism by which local Ca^2+^ signals regulate UPR activation. However, the molecular mechanism linking these Ca^2+^ microdomains to PERK activation remains elusive. Previously, we showed that calcineurin (CN), a Ca^2+^ -dependent heterodimer composed of catalytic (CNA) and regulatory (CNB) subunits, exerts a non-canonical pro-survival function by promoting PERK autophosphorylation. Here, using super-resolution microscopy, CRISPR-Cas9 editing, in silico analyses, and optogenetic droplet assays, we identify CNB as a local Ca^2+^ sensor that couples translocon-generated Ca^2+^ signals to liquid condensate assembly, thereby promoting adaptive PERK phosphorylation. These findings establish CNB-mediated condensate assembly as a mechanism that translates local Ca^2+^ signals into spatially organized early adaptive PERK signaling.

## INTRODUCTION

The ability to detect and respond to cellular stresses, such as the accumulation of misfolded proteins in the endoplasmic reticulum (ER), is essential for maintaining homeostasis in eukaryotic cells (1, 2). A complex signaling network, collectively known as the unfolded protein response (UPR), enables cells to restore homeostasis through the concerted activity of transcriptional and translational regulatory pathways. However, if this cannot be achieved, the UPR itself triggers apoptosis (3). One of the three branches of this cellular response is controlled by PERK (protein kinase RNA [PKR]-like ER kinase), an ER transmembrane protein with a segment that connects a cytosolic domain with kinase activity to a luminal domain whose association with the chaperone BiP (<u>b</u>inding immunoglobulin <u>p</u>rotein) maintains PERK in its monomeric and inactive form. During ER stress, BiP is titrated by the unfolded protein, and PERK is consequently activated by oligomerization and trans-autophosphorylation. Activated PERK phosphorylates the *α-*subunit of eIF2, attenuating global protein synthesis. Importantly, among UPR branches, only PERK remains active during prolonged ER stress (4, 5), thereby mediating both early cytoprotective and chronic pro-apoptotic responses.

Recently, we identified highly localized Ca^2+^ microdomains, particularly during the acute phase of the UPR (6). These Ca^2+^ microdomains are generated by the translocon (7, 8), a core heterotrimeric Sec61 complex (Sec61αβγ) in which the α subunit forms the aqueous pore (9). Sec61α is blocked by the ribosome on the cytosolic side and by BiP on the luminal side (10, 11). During the UPR, phosphorylation of eIF2α prevents recruitment of Met-tRNAi/ribosome to Sec61α, while misfolded proteins sequester BiP, thereby enhancing Ca^2+^ permeability through the Sec61α translocon pore and initiating signaling without classical intracellular Ca^2+^ channels (Inositol triphosphate and Ryanodine Receptors - IP₃Rs and RyRs-). In addition, these local Ca^2+^ events amplify oligomerization of phosphorylated PERK (P-PERK) in cultured human astrocytes, uncovering a Ca^2+^ signaling mechanism whereby stressor-mediated Ca^2+^ increase regulates the UPR (6).

However, the coupling mechanism between P-PERK oligomerization and translocon-mediated Ca^2+^ signaling remains to be determined.

Previously, we showed that calcineurin (CN), a Ca^2+^-dependent protein, promotes cell survival during the acute phase of the UPR through a PERK-dependent mechanism, in both primary mouse and human astrocyte cultures and in two mouse models of brain injury, photothrombotic stroke and traumatic brain injury (TBI) (12, 13).

CN is a heterodimer composed of a catalytic subunit (CNA) and a regulatory subunit (CNB) (14). Two isoforms of CNA are expressed in mammalian brain tissue: α and β (15). The β isoform of CNA rapidly increases post-TBI in astrocytes (13). CNAβ directly interacts with the cytosolic domain of PERK, promoting its autophosphorylation and oligomerization, thereby further reducing protein synthesis. A controlled and localized Ca^2+^ rise enhances this interaction, which requires increased expression levels and is reduced by Ca^2+^–calmodulin (12, 16). This non-canonical cytoprotective function of CN is unrelated to its phosphatase activity, which depends on its binding to Ca^2+^–calmodulin and on exacerbated or sustained Ca^2+^ increases (17).

Taken together, these previous results raise further unanswered questions: how these macromolecules assemble, how they are organized spatiotemporally, and how they sense Ca^2+^. While CNA (α or β) lacks Ca^2+^-binding sites, the regulatory CNB subunit contains two pairs of Ca^2+^-binding EF-hand motifs that form the N- and C-domains. Their defined spatial arrangement may enable high-affinity Ca^2+^ coordination (18, 19). These motifs typically contain a 12-residue segment that coordinates Ca^2+^ through oxygen atoms provided mainly by conserved aspartate (D) residues. A conserved glutamate (E) at position 12 binds the ion in a bidentate manner, completing the coordination geometry, with Ca^2+^-ligand bond distances around 2.3–2.6 Å (not exceeding ∼4 Å) (20).

Here, we sought to determine whether the CNB subunit serves as the Ca^2+^-sensing element within the CN/PERK complex and to define its role in interpreting the elusive Ca^2+^ signals generated by the translocon, including the stress level at which this complex assembles and becomes active during the UPR. To address this, we combined pharmacological, molecular, in silico, and super-resolution imaging approaches. We observed clear Ca^2+^-dependent co-clustering between the endogenously expressed B subunit of CN and PERK, forming spherical, fusion-prone structures that raise new questions regarding their structural, mechanistic, and functional significance.

Over the last decade, the organization of biomacromolecules through liquid-liquid phase separation (LLPS) has emerged as an alternative and dynamic model for the spatial organization of the cell (21, 22). This common mechanism involves macromolecules, such as proteins and nucleic acids, that form reversible liquid-droplet condensates through weak multivalent interactions. This process typically requires the presence of repetitive domains or intrinsically disordered regions (IDRs). Although abundant biochemical and biophysical in vitro evidence supports the existence of these liquid condensates, conclusive evidence for their presence and regulation within cells remains limited.

Here, using an opto-droplet system, we show that CNB/P-PERK co-clusters form dynamic, liquid-like condensates whose behavior is enhanced under ER stress. Together with our findings identifying CNB as the Ca^2+^-sensing component of the complex, these results support a model in which local Ca^2+^ signals promote regulated CNB/P-PERK assembly during the UPR.

## RESULTS

### Mild ER stress induces Ca^2+^-dependent interaction between Calcineurin and PERK

We previously demonstrated that subunit A of CN directly interacts with the cytosolic domain of PERK, promoting PERK autophosphorylation and oligomerization, thereby further reducing protein synthesis (12). This completely novel function of CNA, which exhibits marked cytoprotective effects, is significantly enhanced by moderated Ca^2+^ increases that mimic stress conditions (6, 13).

Here, we first wondered whether this interaction varies with stress severity. To address this question, human astrocyte cultures were stressed with tunicamycin (Tm), an inhibitor of N-linked glycosylation, under different conditions. The interaction between P-PERK and CNA was then assessed by co-immunoprecipitation from an enriched microsomal membrane fraction. A low Tm concentration (0.5 µg/mL) increased the P-PERK/CNA interaction after both short and prolonged incubations. In contrast, a high concentration (2.5 µg/mL), mimicking irreversible stress, resulted in no detectable association (Fig. 1A). Notably, the B subunit of CN (CNB) was also detected as part of this complex under mild, but not severe, stress conditions (Fig. 1B).

**Figure 1:**
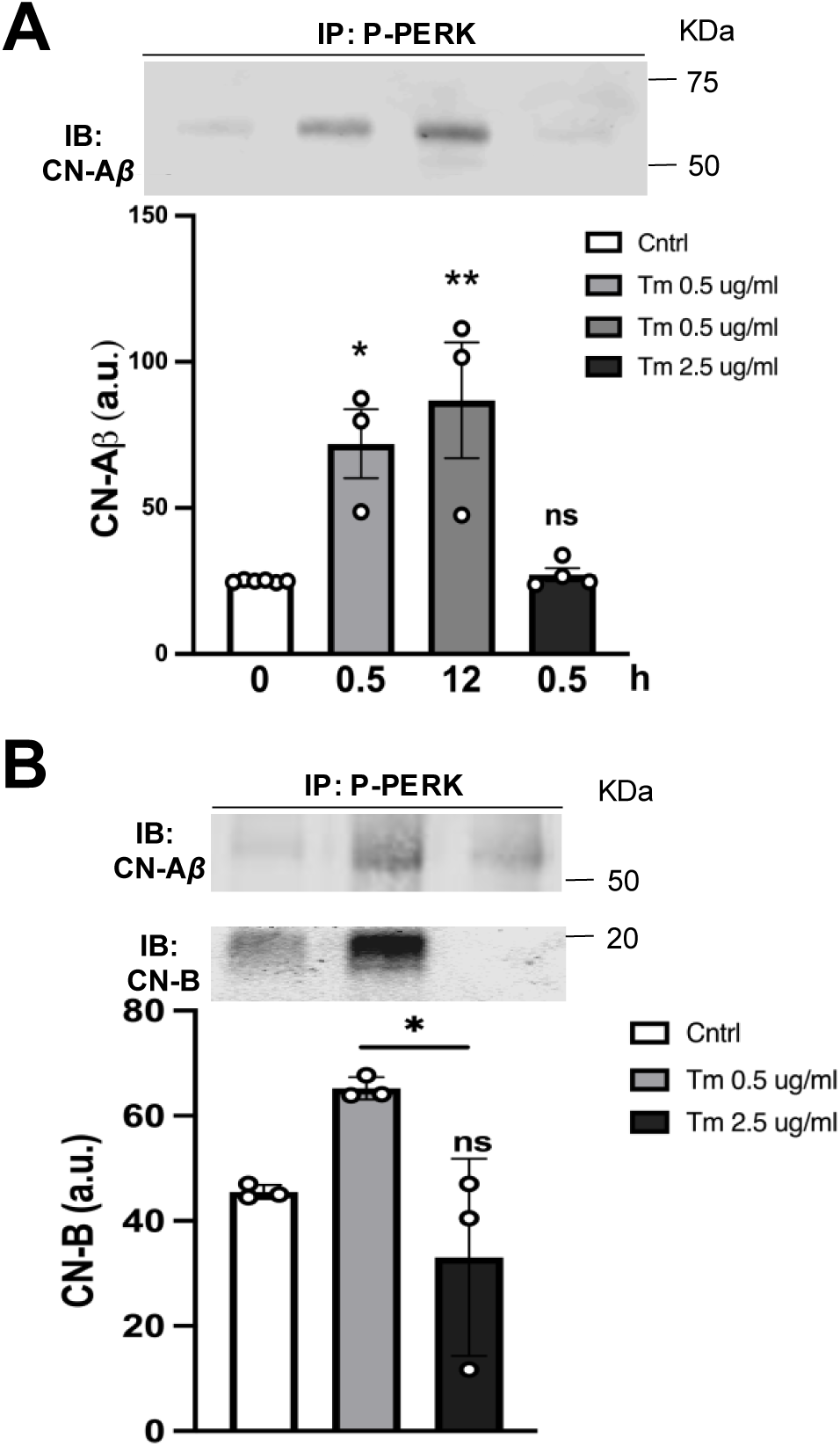
Stress-dependent modulation of the CNAβ/B - PERK interaction. Human astrocyte cultures were treated with two Tm concentrations (0.5 and 2.5 μg/mL) for 0.5 and 12 h. Immunoprecipitation using anti-PERK^UT^ antibody, 12 % SDS-PAGE, and immunoblotting with anti-CNAβ (**A**) and anti-CNB antibodies (**B**). Histograms represent relative intensities (mean ± SEM, ns: no significant difference, *p≤0.05, **p≤0.01). One-way ANOVA test, followed by Tukey’s multiple comparisons, n=3 (**A** and **B**).

Furthermore, to obtain spatial information about the CNB/P-PERK complex, we employed confocal microscopy with an Airyscan detector, achieving a lateral (XY) optical resolution of ∼100 nm. This analysis revealed pronounced co-clustering of the endogenously expressed CNB and P-PERK proteins under mild ER stress conditions (Tm, 0.5 µg/mL for 30 min).

This was initially characterized by line-profile analyses of selected CNB/P-PERK double-positive clusters, revealing a core containing both P-PERK and CNB surrounded by CNB (Fig. 2A). Additionally, co-localization analysis indicates a significant extent of these arrangements at 30 min following Tm treatment (Fig. 2B). Coincident with co-localization, co-cluster size was also significantly increased at this time point. This was determined from the overlap between independently generated green- and red-channel masks, with an average co-cluster size of 0.015 µm^2^ in controls and 0.033 µm^2^ under stress (Fig. 2B).

**Figure 2:**
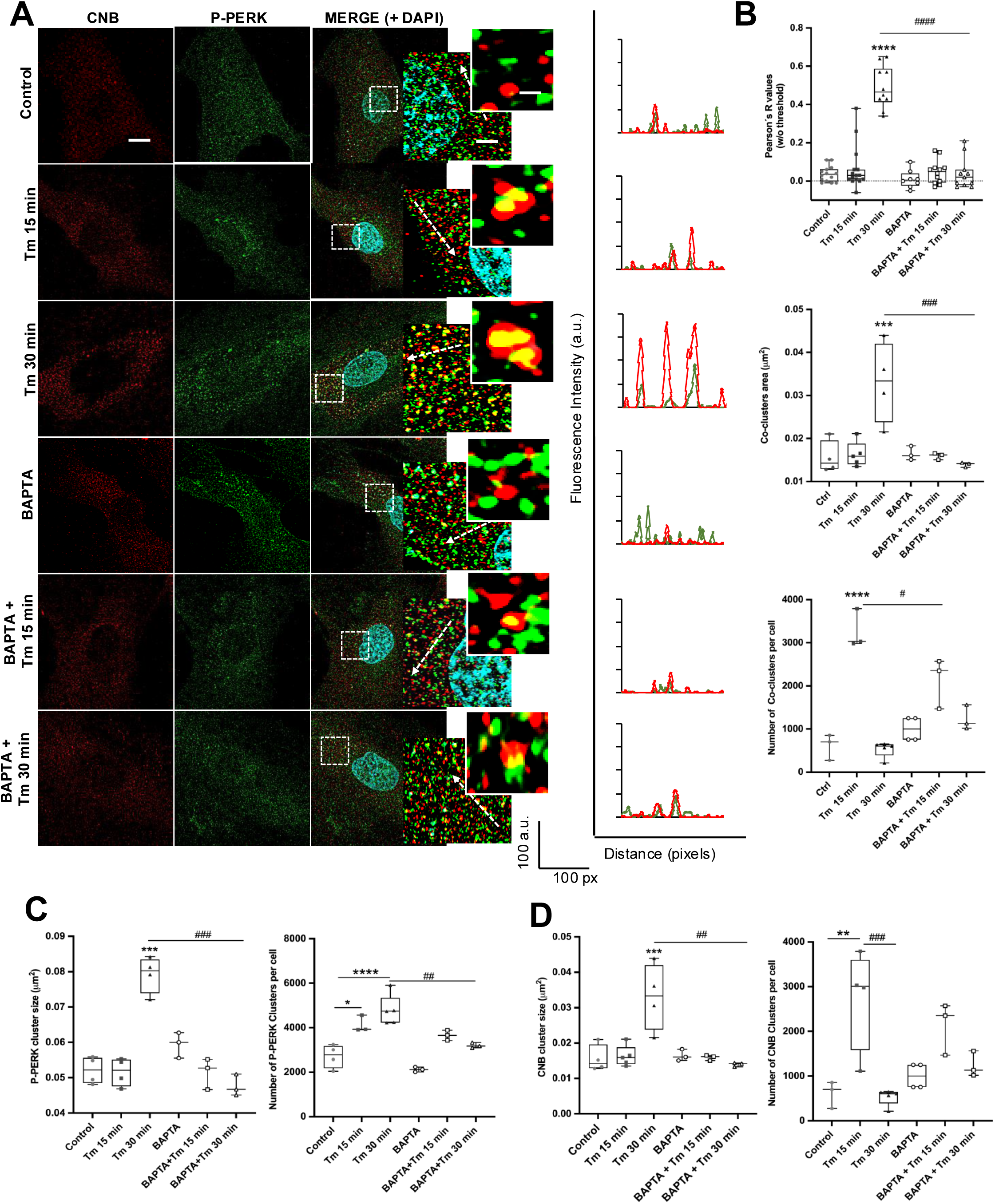
Calcium-dependent P-PERK oligomerization and CNB/P-PERK co-cluster formation in stressed astrocytes. (**A**) Human astrocytes pre-incubated in the absence or presence of BAPTA-AM (20 μM, 30 min), and treated with Tm (0.5 μg/mL for 15 and 30 min). Cells were fixed and immunolabeled with anti-CNB and anti-PERK^UT^ antibodies, followed by Alexa Fluor 568–(red) and Alexa Fluor 488– (green) conjugated secondary antibodies. Nuclei were stained with DAPI (blue). Images were acquired using a Zeiss LSM 980 Airyscan 2 confocal microscope. Scale bars: 11 μm (overview) and 2.75 μm and 0.68 μm (higher-magnification insets). Arrows in each inset indicate the plane with the highest fluorescence signal, used for line-profile generation; the corresponding fluorescence intensity profiles are shown on the right. (**B**) Individual Pearson’s correlation coefficients above the threshold, together with co-clusters area and number, were analyzed. (**C**) P-PERK cluster area and number. (**D**) CNB cluster area and number. (**B**-**D**) Box plots indicate the median and minimum–maximum range. Statistical analysis was performed using one-way ANOVA on mean ± SEM values, followed by multiple comparisons (n = 3; ns, not significant; # and *, p ≤ 0.05; ## p ≤ 0.01; ### and ***, p ≤ 0.001; ****, p ≤ 0.0001).

Comparison of cells exposed to the same concentration of Tm (0.5 µg/mL) for 15 or 30 min revealed a rapid increase in the number of co-clusters per cell within 15 min of treatment. As stress duration increased, co-cluster size increased, whereas the number per cell decreased significantly, suggesting reassembly of the initially formed small complexes. Notably, treatment with BAPTA, a fast Ca^2+^ chelator, significantly reduced changes in all quantified parameters, suggesting that cytosolic Ca^2+^ elevation is required to drive CNB/P-PERK complex assembly (Fig. 2B).

Whereas P-PERK clusters increased in both number and size with increasing stress, the size and number of CNB clusters per cell closely paralleled those of the co-clusters across all stress conditions, suggesting a close association between B clustering and co-cluster dynamics (Fig. 2C, D).

Consistently, co-localization between both CN subunits, A and B, was also increased under stress (Fig. 1S A-B).

To gain further insight into the organization of the CNB/P-PERK complex, we employed stimulated emission depletion (STED) nanoscopy. A key advantage of this super-resolution technique is its ability to simultaneously silence two fluorophores, enabling offset-free colocalization analysis at an optical resolution of approximately 30 nm (23). Compared with conventional confocal microscopy, STED revealed substantially greater structural detail of CNB and P-PERK clusters (Fig. 3A). Compared with our previous super-resolution colocalization analysis (Fig. 2), two-color STED nanoscopy confirmed the close spatial association between CNB and P-PERK and enabled a detailed analysis of the nanoscale organization of the complex. Individual CNB and P-PERK clusters exhibited comparable sizes, ranging from approximately 2,000 to 6,000 nm^2^ (Fig. 3B, C). However, CNB clusters were markedly more abundant than P-PERK clusters, consistent with the proposed core–shell organization of the complex.

**Figure 3:**
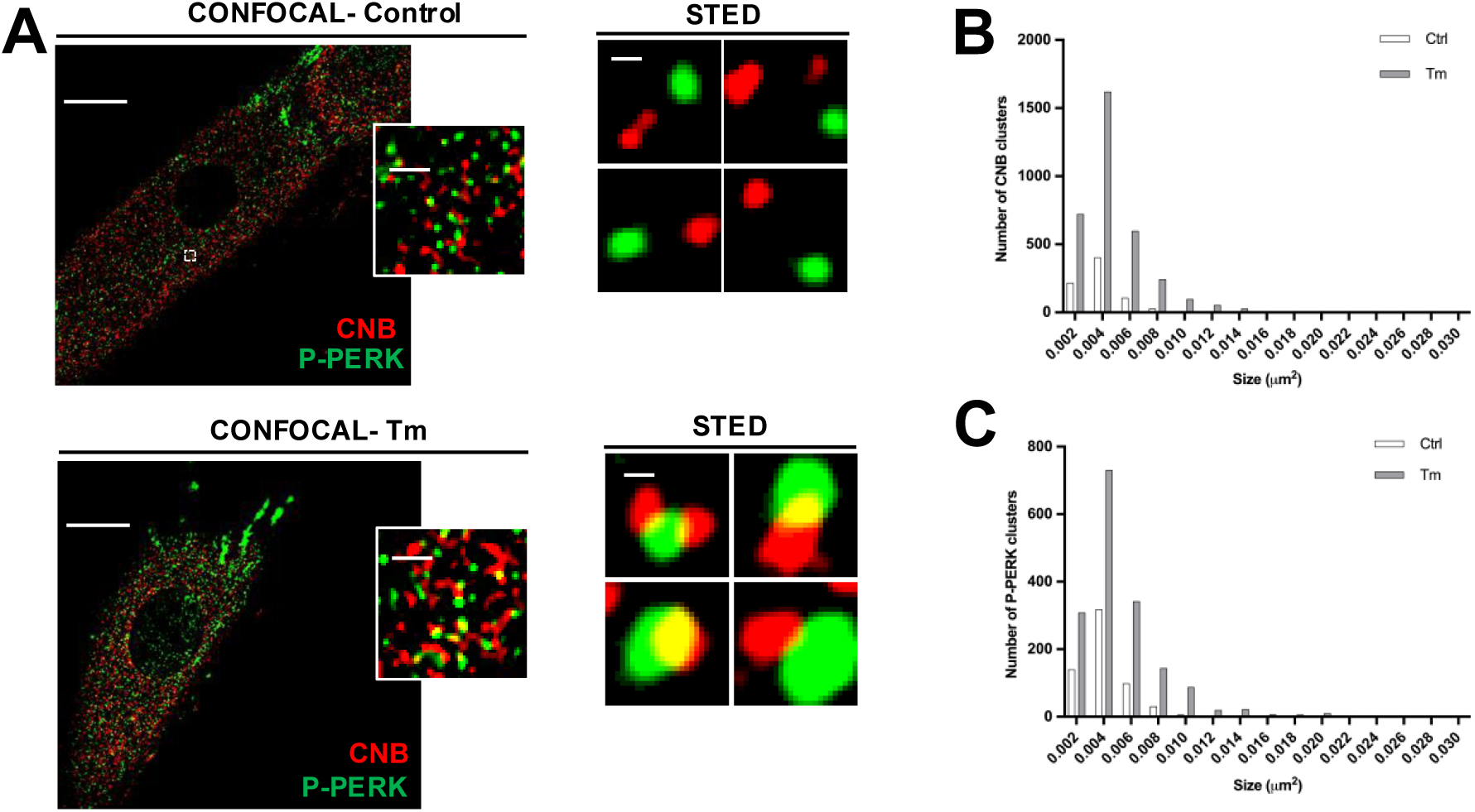
Fluorescence nanoscopy of CNB/P-PERK co-cluster. (**A**) Human astrocytes treated with Tm (0.5 μg/mL for 30 min) were fixed and immunolabeled with anti-CNB and anti-PERK^UT^ antibodies, followed by Abberior STAR RED and STAR ORANGE secondary antibodies. Images were acquired using an Olympus microscope with a STED (stimulated emission depletion) nanoscopy module. Scale bars: 16 μm (confocal images), 0.6 μm (confocal images insets), and 0.06 μm (STED images). (**B, C**) Distribution histograms showing the number of CNB clusters (B) and the number of P-PERK clusters (C) as a function of their as a function of their respective cluster areas.

### Ca^2+^-dependent formation of CNB/P-PERK co-clusters in a cellular model lacking classical intracellular Ca^2+^ channels

To identify the Ca^2+^ source of this signaling, we turned to the use of a Human Embryonic Kidney cell line (HEK-293), in which all three inositol 1,4,5-trisphosphate receptor (IP_3_R) isoforms are knocked out (HEK-TKO) and which also lacks endogenous ryanodine receptors (24). Previously, we used this cellular model to demonstrate stress-induced Ca^2+^ signaling generated by the Sec61α translocon. These cells exhibited highly localized and transient Ca^2+^ microdomains even following treatment with a high concentration of tunicamycin (2.5 µg/mL) (6). Here, Airyscan imaging in HEK-TKO cells revealed Tm-induced CNB/P-PERK co-clusters, which was abolished by BAPTA, suggesting that translocon-induced Ca^2+^ signaling is involved in their formation (Fig. 4A).

**Figure 4:**
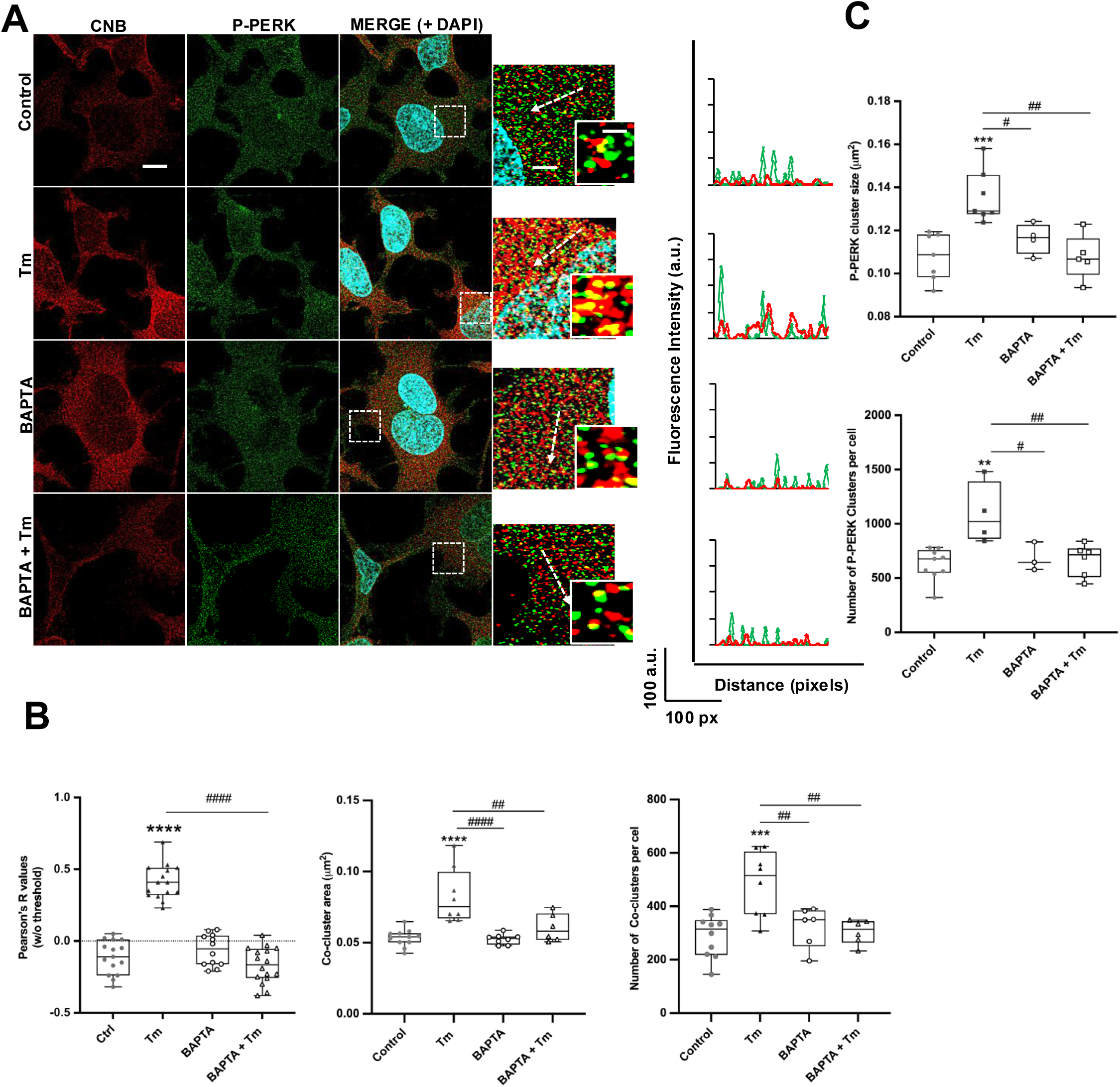
Stress-Induced Ca^2+^ release through translocon promotes CNB/P-PERK co-clusters. (**A**) HEK-TKO cells in low Ca^2+^ buffer were pre-incubated ± BAPTA-AM (20 μM, 30 min) and treated with Tm (0.5 μg/mL, 30 min). Cells were fixed, immunolabeled, imaged, and line profiles generated as in Fig. 2. Scale bars: 11 μm (overview), 2.75 μm and 0.68 μm (insets). (**B**) Quantitative parameters and box plots as in Fig. 2 C. One-way ANOVA on mean ± SEM values, followed by multiple comparisons (n = 3; ns, not significant; ## p ≤ 0.01; ### and ***, p ≤ 0.001; #### and****, p ≤ 0.0001).

Under stress conditions, all parameters used to quantify CNB/P-PERK complex assembly (Fig. 4B), together with P-PERK clusters formation (Fig. 4C), were significantly higher than in the control condition. These parameters were also significantly reduced upon intracellular Ca^2+^ chelation with BAPTA.

In addition, we used pharmacological agents previously described by us to modulate Tm-induced Ca^2+^ microdomains and assessed their effects on PERK product P-eIF2α levels. Specifically, HEK-TKO cells were pre-incubated with anisomycin, a translocon blocker that prevents Tm-induced Ca^2+^ events (6), and then treated with Tm. Confocal imaging revealed that anisomycin significantly attenuated the Tm-induced increase in P-eIF2α levels (Fig. S2A). In contrast, puromycin, which induces premature release of nascent polypeptide chains and enhances translocon-mediated Ca^2+^ signaling, significantly amplified P-eIF2α levels in stressed cells (Fig. S2B).

### The regulatory subunit B as a candidate Ca^2+^ sensor in the Calcineurin/PERK complex

To investigate how this Ca^2+^ signal is detected, we first analyzed the cytosolic portion of PERK using the PROSITE computational service (ExPASy), which revealed no identifiable Ca^2+^-binding sites, including EF-hand motifs. The absence of predicted Ca^2+^-binding sites was corroborated by an *in vitro* kinase assay using a glutathione S-transferase (GST)-fused cytosolic domain of PERK (GST-cPERK), performed at Ca^2+^ concentrations mimicking high (4 µM) and low (50 nM) cytosolic levels (Fig. S3). Under these conditions, PERK autophosphorylation was indistinguishable between Ca^2+^ concentrations.

These findings prompted us to investigate whether CNB instead senses this Ca^2+^ signal. To this end, the CNB-deficient HEK-TKO cell clone was generated using a CRISPR/Cas9 approach by targeting Exon 3 of the PPP3R1 gene (Transgenic Core Facility, UTHSCSA, USA). Such gene editing resulted in a unique cell population carrying a biallelic +1 indel (192-193insA), confirmed by next-generation sequencing, which caused a frameshift leading to a truncated CNB protein with a 64-amino-acid CNB sequence and an additional 8 amino acids before a STOP codon (Fig. S4A, B). The absence of full-length CNB and the expression of truncated CNB were confirmed by immunoblotting (Fig. S4C, D). Hereafter, these cells will be referred to as Truncated-CNB HEK-TKO cells.

First, we assessed the Tm-induced Ca^2+^ increase using a genetically encoded calcium indicator (GECI; GCamP6m) fused to the C-terminal region of cytochrome b5 (Cytb5) to anchor it to the ER membrane (6). Both truncated-CNB HEK-TKO cells and their isogenic control exhibited localized Ca^2+^ events with no significant differences in amplitude (Fig. 5 A, B). However, in cells deficient in full-length CNB, the mean P-PERK cluster size, indicative of kinase activation and oligomerization, does not change significantly under stress or in the presence of BAPTA (Fig. 5C), whereas a significant change is observed in the isogenic control cells (Fig. 4B). This lack of sensitivity to stress and calcium suggests that CNB may act as the Ca^2+^ sensor in this signaling pathway. Indeed, no significant changes were observed in parameters associated with CNB/P-PERK co-cluster formation (Fig. 5D, E). These results indicate that HEK-TKO cells expressing truncated CNB failed to mount any further PERK activation.

**Figure 5:**
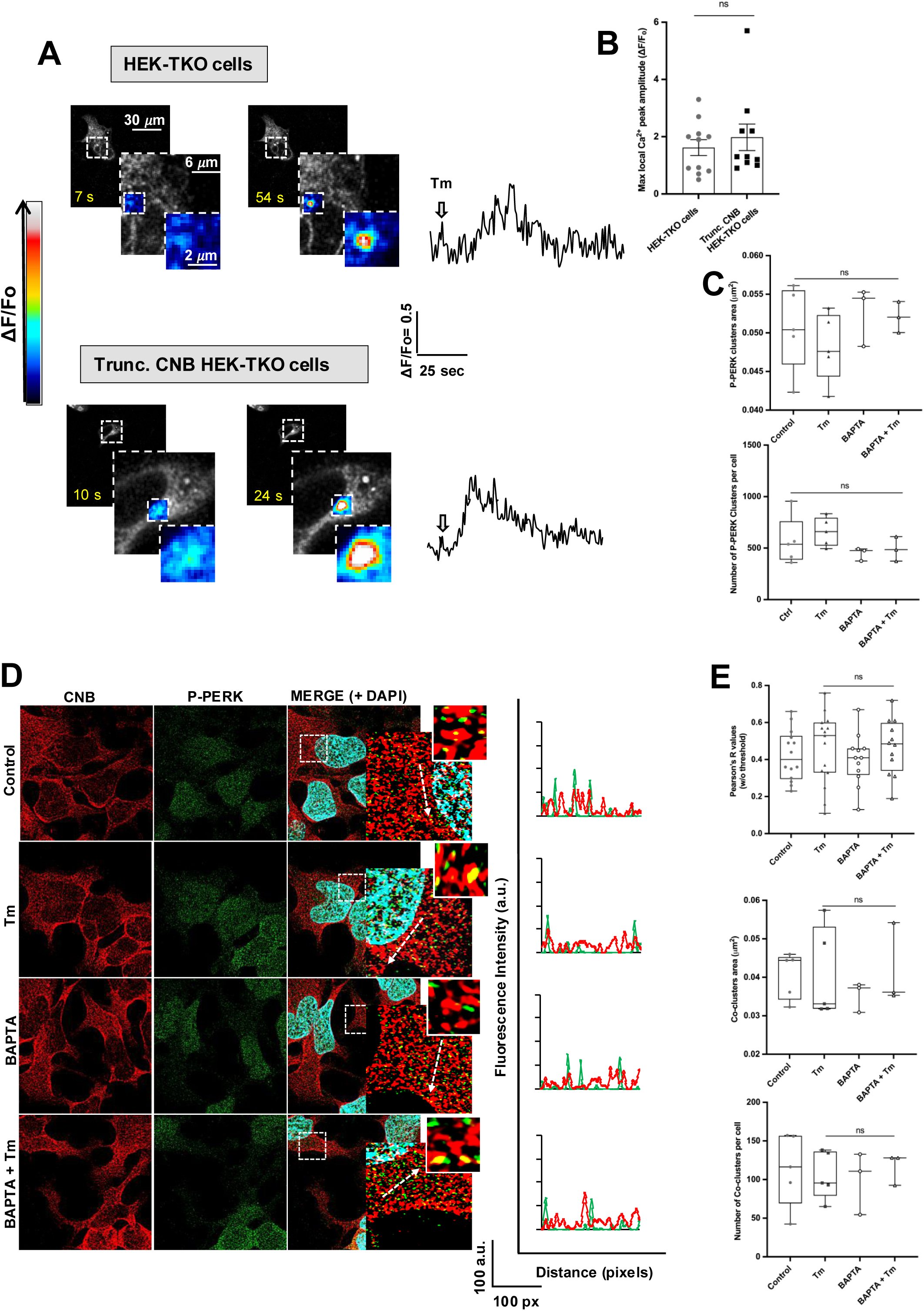
Stress-induced local Ca^2+^ increases are preserved but Ca^2+^ sensing is lost in HEK-TKO cells expressing truncated CNB. (**A**) Measurement of cytosolic Ca^2+^ changes by confocal imaging of cultured HEK-TKO and Trunc. CNB HEK-TKO cells expressing Ca^2+^ indicator GCaMP6-Cytb5. Confocal images corresponding to Ca^2+^ release before and after Tm treatment (Tm; 2.5 μg/mL) at the indicated times. Intensity scale bar for these images is shown. Scale bars: 30 μm (overview), 6 μm, and 2 μm (insets). Fluorescence intensity values were obtained by selecting a 5 x 5-pixel region from subsequent images during the recording of individual astrocytes. Values normalized against values obtained prior to Tm treatment (ΔF/Fo) and plotted as a function of time. (**B**) Histograms (mean ± SEM) showing maximal ΔF/Fo for each condition. (**D**) Cells were fixed, immunolabeled, imaged, and line profiles generated as in Fig. 2. (**C** and **E**) Quantitative parameters and box plots as in Fig 2 B and C. One-way ANOVA on mean ± SEM values, followed by multiple comparisons (n = 3; ns, not significant).

Notably, the truncated CNB variant retains one of the four EF-hand motifs present in the full-length protein. We therefore asked whether this remaining binding site coordinates Ca^2+^ with high affinity (i.e., functions as a structural site) or whether its geometry is distorted, preventing effective Ca^2+^ coordination. To address this, we performed molecular dynamics simulations to examine the relationship among the coordinating residues of the EF-hand closest to the N-terminus. These analyses primarily used ColabFold for structure prediction and PyMOL to model the local positioning of amino acid residues (Fig. 6).

**Figure 6:**
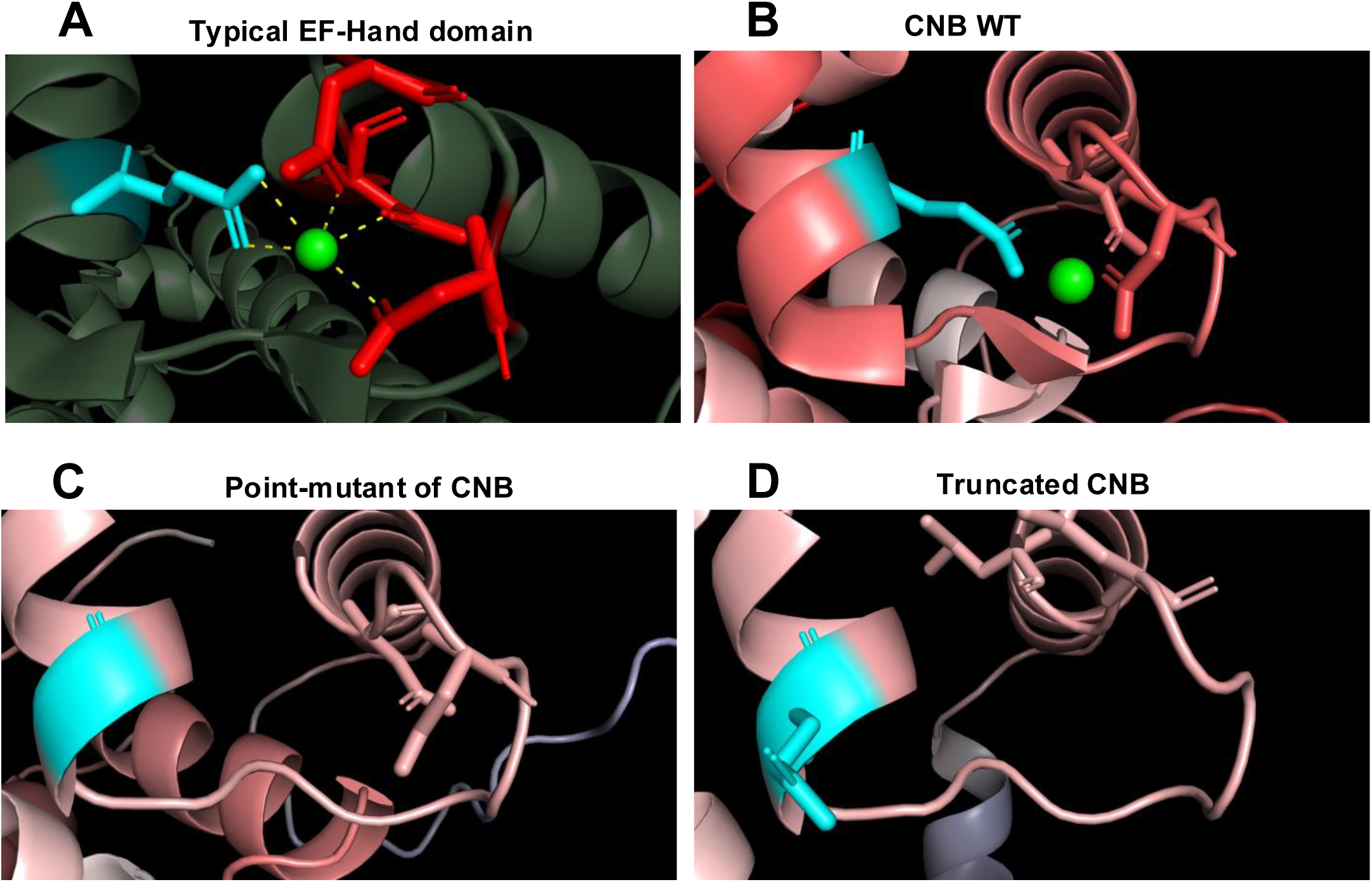
Altered coordination distances at the tentative Ca^2+^ site in Truncated CNB. (**A**) Schematic representation of a typical EF-hand helix–loop–helix motif from N-terminal Calmodulin, showing the position of the coordinated Ca^2+^ ion. Glutamic acid (cyan) and aspartic acid (red) residues are highlighted in stick representation for clarity, alongside a cartoon representation colored by the protein’s B-factor values and the Ca^2+^ ion (green sphere). Note the six coordinating amino acid residues located within the loop at positions 1, 3, 5, 7, 9, and 12, with the glutamate at position 12 binding the Ca^2+^ in a bidentate manner and completing the coordination geometry. (**B**-**D**) The N-terminal EF-hand motif of CNB WT, PDB *ID* 4OR9 (**B**), the point mutant CNB_E41, E74, E111, E151_ (**C**), and the truncated CNB variant (**D**) are shown. Note that the Ca^2+^ ion coordinated in CNB WT is absent in the CNB mutant because of the lack of a coordinating oxygen at position 12, and in the truncated CNB variant because of the distortion of the loop geometry. (**A**-**D**) The illustrations were generated in PyMOL v3.1.6.1.

The first simulation modeled Ca^2+^ coordination in the wild-type CNB subunit and in a point-mutant CNB variant in which the four conserved glutamate residues (E41, E74, E111, and E151) were replaced with glycine (E→G), serving as a negative control. This mutation (E41→G) caused a clear distortion of the Ca^2+^-binding site geometry, moving the oxygen ligands of aspartate residues (D30 and D32) to distances greater than 4 Å from the Ca^2+^ ion, thereby preventing effective ion coordination. In a second simulation using the truncated CNB variant, which retains the conserved E41 residue, an inappropriate orientation of this side chain was observed. This prevented both bidentate and monodentate coordination and was consistent with the experimentally observed Ca^2+^ insensitivity (Fig. 6).

### ER stress enhances CNB condensate formation

To investigate whether CNB/PERK co-clustering could involve liquid-liquid phase separation (LLPS), we examined the presence of intrinsically disordered regions (IDRs) in CNB and PERK, as well as in CNA, a protein shown to associate with the CNB/PERK signaling complex (Fig.1 and (12)). Analysis using the Predictor of Natural Disordered Regions (PONDR VL-XT; (25)) revealed predicted IDRs in all three proteins (Fig. S5A-C). In particular, PERK exhibited extensive disorder throughout its sequence, including the juxtamembrane (JM) and cytosolic domains, which contain eight unstructured segments enriched in serine (S), proline (P), and asparagine (N) residues. By contrast, CNB displayed a single disordered N-terminal region enriched in charged residues (K, R, D, and E), a sequence signature characteristic of IDRs.

Notably, the truncated CNB variant also contains predicted IDRs (residues 1–10 and 17– 50) (Fig. S5D), raising the possibility that it may retain the ability to undergo constitutive CNB/P-PERK co-clustering (Fig. 5D, E), whereas the lack of functional EF-hand domains suggests that these domains may mediate the Ca²⁺-dependent enhancement of CNB/P-PERK co-clustering during ER stress.

Motivated by the presence of predicted IDRs in CNB, we employed an optogenetic system (26) to investigate whether CNB undergoes liquid-like condensation in the absence or presence of ER stress. HEK-TKO cells expressing either mCherry-CRY2must alone or CNB fused to mCherry-CRY2must (mCh-CRY2must-CNBwt) were exposed to a 10-sec pulse of blue light (Fig. 7). After photoactivation under control conditions, both mCh-CRY2must and mCh-CRY2must-CNBwt rapidly formed numerous small light-induced clusters.

**Figure 7:**
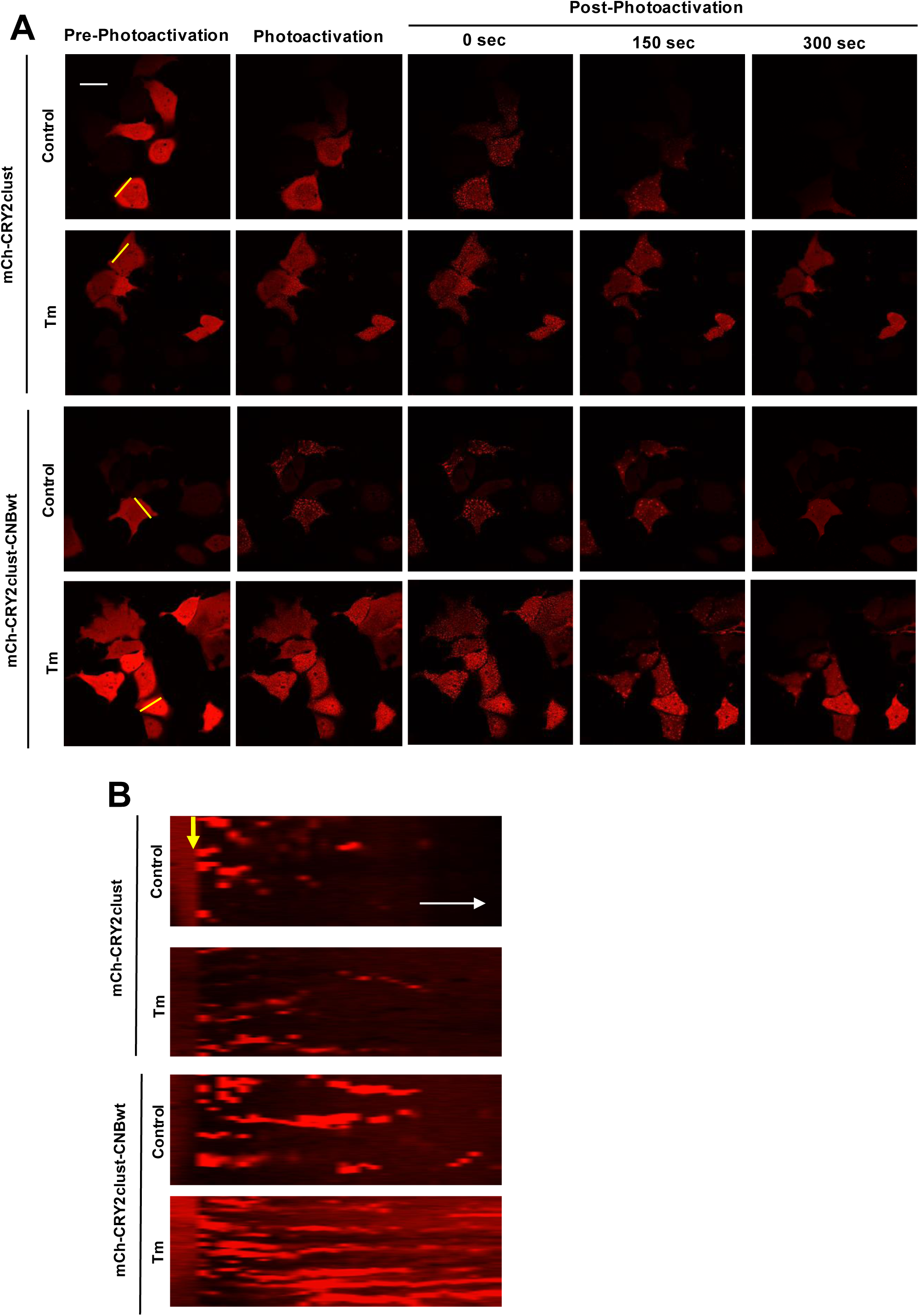
Optogenetic induction of CNB condensates. HEK-TKO cells expressing either mCh-CRY2must alone or mCh-CRY2must-BBwt were imaged using a Zeiss LSM 980 Airyscan 2 confocal microscope. (**A**) Representative pseudocolored images acquired before and after photoactivation with a 10 sec pulse of blue light at the indicated time points. Tunicamycin (Tm, 2.5 μg/mL) was added at the onset of photoactivation. (**B**) Kymographs generated from the yellow line scans shown in A). Yellow arrows indicate the time of photoactivation. The white arrow indicates the direction of time and the corresponding time interval.

Notably, the assembly of mCh-CRY2must-CNBwt condensates was significantly enhanced when light induction was combined with Tm treatment, consistent with a role for ER stress-induced local Ca^2+^ signaling in this process. Representative kymographs (Fig. 7B) illustrate the dynamic behavior of the optogenetically induced condensates. Under all conditions, condensates formed immediately after photoactivation and exhibited directional movement over time. However, CNB condensates persisted longer than controls and, under ER stress conditions, were markedly more abundant and frequently underwent fusion and fission events, behaviors consistent with liquid-like condensates.

Taken together, these findings indicate that the CRY2 optodroplet module provides the initial nucleation trigger, whereas CNB, likely through its intrinsically disordered regions, promotes the stabilization and persistence of the resulting condensates. ER stress further potentiates this process, consistent with the idea that stress-induced changes in the intracellular microenvironment facilitate CNB condensation.

## DISCUSSION

In this study, we demonstrate that subunit B of CN is required for the signaling pathway linking Ca^2+^ release across the translocon to PERK phosphorylation and subsequent kinase activation. This process, which primarily occurs under mild and reversible ER stress, involves the formation of dynamic co-clusters between CNB and P-PERK in a Ca^2+^-dependent manner. Indeed, BAPTA markedly reduced the formation of P-PERK clusters, CNB clusters, and CNB/P-PERK co-clusters in both human astrocytes and HEK-TKO cells. Furthermore, experiments using HEK-TKO cells expressing truncated CNB, together with in silico analyses, indicate that Ca^2+^ is sensed by the EF-hands of CNB, promoting conformational changes within the subunit and enhancing CNB-PERK binding. These results, observed in both human astrocytes and HEK-TKO cells, suggest that the assembly of PERK clusters with CNB is not limited to a specific cellular context but also represents a cell-type-independent event.

Super-resolution microscopy demonstrated that higher-order oligomerization of PERK serves as the initiating trigger and constitutes the condensate core, whereas CNB is enriched at the periphery. This spatial organization was revealed by super-resolution imaging at a scale substantially finer than the dimensions of the condensates. Importantly, these condensates were composed of endogenous proteins, strengthening the physiological relevance of these observations and minimizing the likelihood of overexpression-related artifacts. Moreover, our findings provide mechanistic insight into the physicochemical nature of these condensates by demonstrating that they undergo ER membrane-associated liquid–liquid phase separation (LLPS). The first indication of this behavior comes from in silico analysis using PONDR VL-XT, which predicts extensive intrinsic disorder in both proteins, a structural feature known to promote the weak multivalent interactions that drive LLPS.

In addition, we directly assessed the phase separation behavior of CNB using an optogenetic approach. CNB fused to the CRY2 must module readily formed condensates upon blue-light stimulation and tunicamycin treatment. These optogenetically induced condensates closely resembled the spherical morphology of endogenous CNB/PERK co-clusters. Moreover, the time-dependent remodeling of endogenous CNB/PERK co-clusters observed in fixed cells— characterized by an early increase in cluster number followed by fewer, larger condensates after prolonged tunicamycin exposure—is consistent with the maturation and coalescence of initially formed condensates. Consistent with this interpretation, optogenetically induced CNB condensates exhibited hallmark liquid-like behaviors, including fusion, dynamic mobility, and reversibility.

Notably, the truncated CNB variant also contains intrinsically disordered regions (IDRs; residues 1–10 and 17–50), which could explain its ability to form condensates under basal conditions. However, the absence of functional EF-hand domains suggests that these motifs are required to sense stress-associated Ca^2+^ signals and regulate phase separation in response to mild stress. Although cytosolic Ca^2+^ has been implicated in regulating other liquid condensate systems (27, 28), the present study provides molecular insight into how local Ca^2+^ microdomains are sensed by EF-hand domains. Moreover, the loss of these motifs leaves cells with a constitutively active UPR, supporting the notion that EF-hand domains function as a molecular switch that controls cellular stress responses.

This non-canonical function of CNB, acting in concert with the CN-Aβ catalytic isoform, is PERK-dependent and mechanistically distinct from the classical phosphatase activity of calcineurin, which has been linked to neuronal death (17). Instead, the CN-Aβ-CNB–PERK signaling axis promotes cellular adaptation and survival under stress by inducing controlled increases in intracellular Ca^2+^ levels (12, 13, 29).

Importantly, this function depends on stress-induced upregulation of both CNAβ and CNB, as observed in vitro and in vivo (12, 13). These findings are particularly relevant given a naturally occurring single-nucleotide polymorphism in PPP3R1 (rs1868402), the gene encoding the regulatory B subunit of calcineurin, which is associated with reduced CNB expression and accelerated progression of Alzheimer’s disease (30). Although the underlying mechanism remains unknown, this genetic association raises the possibility that reduced CNB expression may compromise the non-canonical function of the CNAβ–CNB heterodimer in coupling ER stress-induced Ca^2+^ signals to the acute, cytoprotective regulation of PERK signaling. Such a mechanism may help preserve early PERK signaling while limiting the transition to chronic PERK activation associated with neurodegeneration.

Finally, our findings reveal that CNB-mediated Ca^2+^ sensing is coupled to the assembly of liquid condensates, providing a mechanism by which local Ca^2+^ signals are translated into spatially organized, adaptive PERK signaling. This work provides a conceptual link among liquid-phase condensation, subcellular organization, and the restoration of cellular homeostasis during the early response to ER stress.

Future studies will focus on identifying additional macromolecules that may be recruited to the CNB/P-PERK complex, forming a dynamic signaling platform that coordinates its assembly, dynamics, and function.

## MATERIALS AND METHODS

### Reagents, antibodies and constructs

The mouse cDNA encoding CNAβ (isoform β of CN-A) followed by mouse (Mus musculus) cDNA encoding CNB (CNAβ-CNB) was subcloned into the mammalian vector mCherry-CRY2clust (Addgene, Plasmid #105624), and it was termed **p**mCherry-CRY2clust/CNAβ-CNB. The genetically encoded calcium indicator (GECI) GCaMP6m cDNA was subcloned into the pcDNA3.1 vector, which was engineered to include the cDNA encoding the C-terminal 76 amino acid residues of rat cytochrome b5, as described in (6). The resulting construct was termed **p**GCaMP6m-Cytb5.

Reagents were from Fisher Scientific or Sigma-Aldrich unless specified otherwise. The primary antibodies we used in this study were: in-house anti-PERK^UT^ antibody (12), which is now commercially available from Biosensis (Australia-USA, # R-1333-100); anti-CN-Aβ (#07-068-I), (Sigma-Aldrich); anti-CNB (# MA5-23933) (Invitrogen).

### Cell culture and transfection

Human astrocytes were obtained from different donors (1 male patient aged 56 years and 3 female patients aged 36-50 years) sourced from brain biopsies of anonymous patient waste tissues and purified from non-infiltrated tumor brain regions at least 2 cm away from the contrast-enhancing tumor core or from the entry cortex of epilepsy surgeries. The protocol was received approval from the Institutional Ethics Committee of Hospital Privado Universitario de Córdoba, Argentina. Human astrocytes were cultured essentially as described in (16, 31). Briefly, brain tissues were minced using a sterile razor and trypsinized (trypsin-EDTA 0.25%) for 30 min in a 37 °C humidified incubator. Cells were suspended in fresh DMEM/F-12 (#11,039– 021) supplemented with 10% FBS (#12483–020), 10,000 U/ml penicillin, and 10 mg/ml streptomycin (#15140–122).

Triple IP3R knockout human embryonic kidney cell line (TKO-HEK) from Kerafast (Boston, MA, USA; #EUR030), as well as the truncated-CNB TKO-HEK cells, were grown in DMEM (#11995–065) supplemented with 10% FBS, 10,000 U/ml penicillin, and 10 mg/ml streptomycin. All cell cultures were incubated at 37 °C in a humid 5% CO_2_ atmosphere. Cell cultures were tested for the presence of Mycoplasma using a PCR-based method.

TKO-HEK and truncated-CNB TKO-HEK cells were plated in either 35 mm dishes (#P35G-1.5–14-C, MatTek Corp.; Ashland, MA, USA) as indicated, and transfected with either 2 μg cDNA pGCamP6m-Cytb5 and 2 μl transfection agent X-tremeGENE (#06366244001), according to the manufacturer’s instructions. Calcium imaging was performed 2 days after transfection in both cell types.

TKO-HEK cells were transfected with either **p**mCherry-CRY2clust/CNAβ-CNB or an empty vector, using the same μg cDNA/μL X-tremeGENE ratio. Optodroplet assay was performed 2 days after transfection.

### Co-immunoprecipitation and Western blotting

Treated cultured cells were washed twice with PBS and lysed in cell lysis buffer (15 mM Tris-HCl, pH 7.6, 140 mM NaCl, 250 mM sucrose, 1 mM EDTA) containing protease and phosphatase inhibitor cocktails.

Co-immunoprecipitation was essentially performed as previously described (12), except that the anti-PERK antibody was first crosslinked to A/G PLUS agarose beads (# sc-2003, Santa Cruz Biotechnology) using disuccinimidyl suberate (DSS) overnight at 4 °C on a rocking platform (32). Each microsome-enriched fraction was then incubated with 40 μl of the antibody-bead complexes for 1 h at room temperature. Immunoprecipitated proteins were eluted from the cross-linked antibody using Elution Buffer (100 mM glycine, pH 2.8, 1% Nonidet P-40). Subsequently, sample buffer supplemented with β-mercaptoethanol was added, and samples were heated at 100 °C for 5 min, followed by centrifugation at 600 × g for 1 min at room temperature, resolved on 7% SDS-PAGE, transferred to a nitrocellulose membrane, and detected by immunoblotting using either CN-Aβ or CNB antibodies.

For Western blotting, proteins from the cytosol-enriched fraction (12) were resolved either by 12% SDS-PAGE and transferred to nitrocellulose membranes or, for detection of low-molecular-weight peptides, by 16% SDS-PAGE and transferred to PVDF membranes. The PVDF membranes were then fixed with 5% glutaraldehyde for 30 min. Both membrane types were analyzed by immunoblotting with an anti-CNB antibody.

All immunoreactive bands generated in Western blot and co-immunoprecipitation assays were visualized using the Odyssey® infrared imaging system.

### *In vitro* Kinase assay

The autophosphorylation of cPERK was performed essentially as described in (12). Briefly, the GST-cPERK, at different concentrations, was incubated for 30 min at 30 °C in 30 µl of reaction mixture and 0.1 mM ATP, 50 µCi [γ^32^P]ATP (6,000 Ci/ mmol, PerkinElmer Life Sciences, Inc., USA) in the presence of two free Ca^2+^ concentrations (50 nM or 4 µM, according to Max Chelator, developed at Stanford University, USA). The reactions were stopped, and the proteins were resolved through 10% SDS-PAGE. The gel was fixed, dried, and autoradiographed to visualize proteins.

### Cell Line and Genetic Editing

CNB-deficient HEK-TKO clones were genetically edited using CRISPR/Cas9 technology. Three clones were generated using the following sgRNAs: 5′-AAATGAGGCTTATCCTTTGG-3′ (exon 2), 5′-TCTATTACTCGCTGTACTAAAGG-3′ (exon 3), and 5′-TAGATATATTCGACACAGATGGG-3′ (exon 3). These clones were validated by Western blotting at the Mouse Genome Engineering and Transgenic Facility (UTHSCSA, USA). Subsequently, the indel mutation in clone #3 was confirmed by next-generation sequencing (NGS) (Macrogen, South Korea).

### Immunocytochemistry

For immunofluorescence detection, cells were grown on 12-mm glass coverslips, washed twice with PBS, fixed with 4% paraformaldehyde and 120 mM sucrose in PBS for 30 min at 37 °C, permeabilized for 5 min with 0.01% Tween 20 and 0.01% digitonin in PBS, blocked for 45 min in 5% BSA in PBS, and incubated overnight at 4 °C with anti-PERK^UT^ (1:400), anti-CNB (1:400), or anti-CN-Aβ (1:400) antibodies as is indicated, and diluted to the indicated concentration in 5% BSA in PBS.

Images were acquired using either a Zeiss LSM 980 equipped with Airyscan 2 and a 60 x oil-immersion objective (NA 1.4), or an Olympus IX81 microscope equipped with an Abberior STEDYCON (stimulated emission depletion) super-resolution module and a UPLXAPO 100 x /1.45 NA oil-immersion objective (CEMINCO, UNC). STED images were acquired with a 10 nm pixel size, 10 µs pixel dwell time, 5 line accumulations, and a 60 µm pinhole. Fluorophores were excited at 561 nm (Abberior STAR ORANGE, item # STORANGE) or 640 nm (Abberior STAR RED, item # STRED). Both fluorophores were depleted using a single 775 nm STED laser.

### Ca^2+^ imaging

Cytosolic Ca^2+^ imaging was performed as previously described (6). Briefly, HEK-TKO and Trunc. CNB HEK-TKO cells were transfected with 2 μg of pGCaMP6m-Cytb5 cDNA and 2 μl of the transfection reagent X-tremeGENE (Sigma-Aldrich). Ca^2+^ imaging was carried out 48 hs later using a Zeiss LSM 800 confocal microscope (CEMINCO, UNC) equipped with a 60 x oil-immersion objective (NA 1.4), with excitation and emission wavelengths at 488 nm and 509 nm, respectively. Prior to imaging, the culture medium was replaced with a low-Ca^2+^ buffer. Images were acquired at 1 s intervals for 5 min, and Tunicamycin (Tm) was added 20 s after the start of recording.

### OptoDroplet assay in living cells

OptoDroplet assays were performed using the empty vector mCherry-CRY2clust (Addgene plasmid #105624). In addition was generated: mCherry-CRY2clust-CNAβ-CNB. HEK TKO cells were transfected as previously described, and experiments were conducted 48 h after transfection.

Live-cell imaging was performed using the 561 nm channel. Basal fluorescence was recorded for 1 min prior to stimulation, acquiring images every 20 s. OptoDroplet formation was then induced by illumination with a 488 nm laser (21 mW) for 10 s. Following stimulation, time-lapse imaging was continued for 5 min with images acquired every 20 s.

### Imaging analysis

All acquired images were deconvolved using TRUESHARP online deconvolution (Version 1.0; Abberior Instruments GmbH, Göttingen, Germany; https://app.truesharp.rocks/). Line profiles of fluorescent signals were generated using ImageJ (Fiji). Images were first split into two channels, and a line was drawn using the “Straight” tool. The mean pixel intensity at each position (xᵢ) along the line was then calculated

For co-localization assays, Pearson’s correlation coefficients were calculated within 5 x 5-pixel ROIs selected from regions showing the most evident fluorescence signal.

The number and area of either the P-PERK clusters or the CNB/P-PERK and CNB/CNA co-clusters per cell were analyzed using the ImageJ plug-in Analyze Particles, by generating a mask using the Otsu Threshold.

For Ca^2+^ imaging analysis, fluorescence intensity values were plotted as ratios (ΔF/Fo) of change of fluorescence (ΔF) of the ROI (5 x 5 pixels) divided by mean resting fluorescence (Fo) prior to Tm addition vs. recording time. ROIs were defined as active when fluorescence increased ≥2 SD relative to baseline fluorescence.

For opto-droplet experiments, kymographs were generated by drawing line regions of interest (ROIs) and applying the Reslice function (Image → Stacks → Reslice) with an output spacing of 1 pixel.

### In silico studies

The CNB wild-type (WT) starting model coordinates for each simulation were derived from the 2.23 Å atomic-resolution *homo sapiens* CNA-CNB cryo-EM structure CNB (PDB, accession code ID: 4OR9) (33). Subsequently, using ColabFold (https://github.com/sokrypton/ColabFold), five structural models were generated for each of the two variants: the point-mutant CNB and the truncated CNB, without using structural templates from PDB ID: 4OR9. For each variant, the model with the highest predicted local distance difference test (pLDDT) score was selected for further analysis and compared with the available cryo-EM structure of CNB WT (PDB ID: 4OR9).

All structural models were visualized using PyMOL (v3.1.6.1). Distances between the aspartate and glutamate residues present in the EF-hand loop motif and the Ca^2+^ ion were subsequently measured in PyMOL. For the point-mutant CNB and truncated CNB variants, the Ca^2+^ ion was manually positioned prior to distance measurements, as it was absent from the predicted structural models.

### Statistical analysis

Statistical analyses were performed using two-sided unpaired t-tests and one-way or two-way analysis of variance (ANOVA), followed by the appropriate post hoc test for multiple comparisons, using GraphPad Prism 10.4.1. Results are presented as mean ± SEM of 3 or more independent replicates. Differences with p-values ≤ 0.05, ≤0.01, ≤0.001 and ≤0.0001 are indicated.

## Acknowledgements

The authors thank Dr. Laura Montroull for cell culture technical support. This study was supported by grants from: National Institutes of Health (NIH), USA (#RO1AG058778-01A1; Subaward Agreement No. 165148/165147 between UTHSCSA-Instituto Investigación Médica M y M Ferreyra), and Agencia Nacional de Scientific and Technological Promotion, Argentina (ANPCyT, PICT 2019 #00155).

Macarena Fernandez were supported by fellowships from The National Scientific and Technical Research Council (CONICET), Argentina.

## Author contributions

Conceptualization: M.B.♦. Experiments: S.B., M.F., G.Q., A.P., D.H, J.C.P., S.A. Data analysis: S.B. Visualization: S.B. MB, M.B.♦. Manuscript writing: S.B., M.B.♦. Supervision: M.B., M.B.♦. Writing – Review & Editing: S.B. M.M., G.E.G., M.B., M.B♦. Funding acquisition: J.D.L., M.B.♦ Project administration: M.B.♦

