## Supplemental data -Figures and Fig. legends for "Calcineurin B-mediated Ca^2+^ sensing translates stress signal intensity into the assembly of phase-separated condensates at PERK complexes"

**Figure S1. CNB and CNA show increased colocalization under stress conditions.**
(**A**) Human astrocytes treated with Tm (0.5 μg/mL, 30 min) were fixed and immunolabeled with anti-CNB and anti-CNA antibodies, followed by Alexa Fluor 568–(red) and Alexa Fluor 488–(green)–conjugated secondary antibodies. Nuclei were stained with DAPI (blue). Images were acquired using a Zeiss LSM 980 Airyscan 2 confocal microscope. Scale bars: 22 μm (overview) and 1.1 μm (inset). (**B**) Individual Pearson's correlation coefficients above the threshold were analyzed. Box plots indicate the median and minimum–maximum range. Statistical analysis was performed using one-way ANOVA on mean ± SEM values, followed by multiple comparisons (n = 3; ****, $p \le 0.0001$).

**Figure S2: Modulation of translocon-mediated $Ca^{2+}$ release alters tunicamycin-induced eIF2α phosphorylation.** TKO-HEK cells were pre-incubated with or without anisomycin (Aniso, 100 μM, 30 min) followed by treatment with tunicamycin (Tm; 0.5 or 2.5 μg/mL, 30 min) (**A**), or pre-treated with puromycin (Puro, 200 μM, 5 min) prior to exposure to Tm (0.5 μg/mL, 30 min) (**B**). Cells were then fixed and immunolabeled with an anti-P-eIF2α primary antibody and an Alexa Fluor 488–conjugated secondary antibody (green). Nuclei were visualized with DAPI (blue). (**B**) In addition to P-eIF2α (green) and DAPI (blue), F-actin was labeled with rhodamine-phalloidin (red). Images in (**A**–**B**) were acquired using a Zeiss LSM 800 confocal microscope. Scale bar: 60 μm. Histograms (mean ± SEM) show either P-eIF2α fluorescence intensity (**A**) or Manders' overlap coefficient (**B**). Statistical analysis was performed using one-way ANOVA followed by multiple comparisons (n = 3; *, $p \le 0.05$; ## $p \le 0.01$; ***, $p \le 0.001$).

**Figure S3: PERK Autophosphorylation Is Independent of $Ca^{2+}$ Levels**

Recombinant GST-cPERK was incubated with [γ-$^{32}$P]ATP; a schematic of the autophosphorylation assay is shown in (**A**). (**B**) GST-cPERK at different concentrations was incubated under low (50 nM) and high (3.2 μM) $Ca^{2+}$ conditions in the presence of [γ-$^{32}$P]ATP. Samples were resolved by 10% SDS-PAGE and visualized by autoradiography, as described in Materials and Methods.

**Figure S4: Genetic editing of CNB and the expected protein product.**

(**A**) Identification and quantification of the prevalence of the indel in the genetic editing clone (#3) compared to the isogenic control clone by next-generation sequencing. (**B**) Representative view of the expected amino acid sequence of the full-length CNB from the truncated version (clone #3 product), which retained the first 64 residues and amino acids, and due to the frameshift mutation, an additional 8 amino acids (in green); the presence of the STOP codon is highlighted by the asterisk. Representative CNB immunoblots showing the full-length protein in HEK-TKO whole-cell lysates (**C**) and the truncated form in Trunc. CNB HEK-TKO cells (**D**).

**Figure S5: Prediction of intrinsically disordered regions in CNB, PERK, CNA, and truncated CNB**

Predicted intrinsic disorder profiles of CNB (**A**), PERK (**B**), CNA (**C**), and truncated CNB (**D**), generated using the PONDR VLXT algorithm.

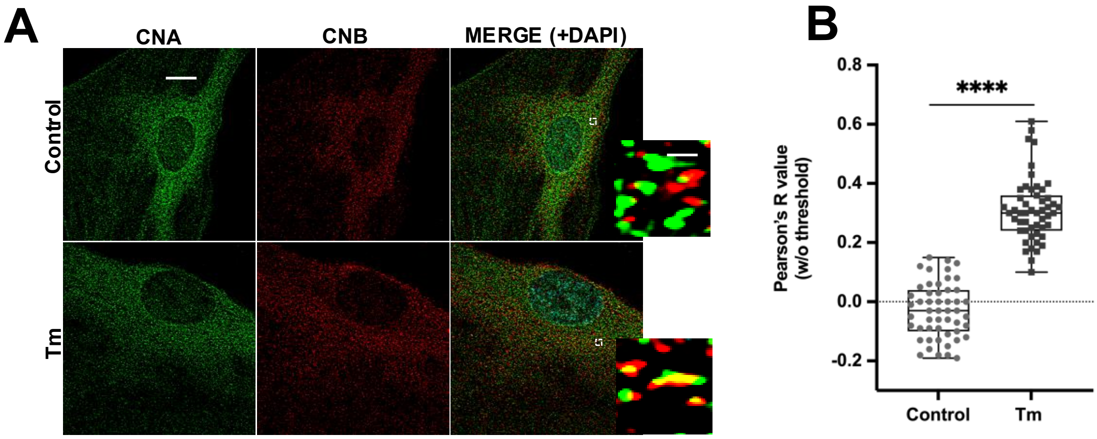
A
CNA
CNB
MERGE (+DAPI)
Control
Tm
B
0.8
0.6
0.4
0.2
0.0
-0.2
****
Pearson's R value
(w/o threshold)
Control
Tm

Figure S1

A

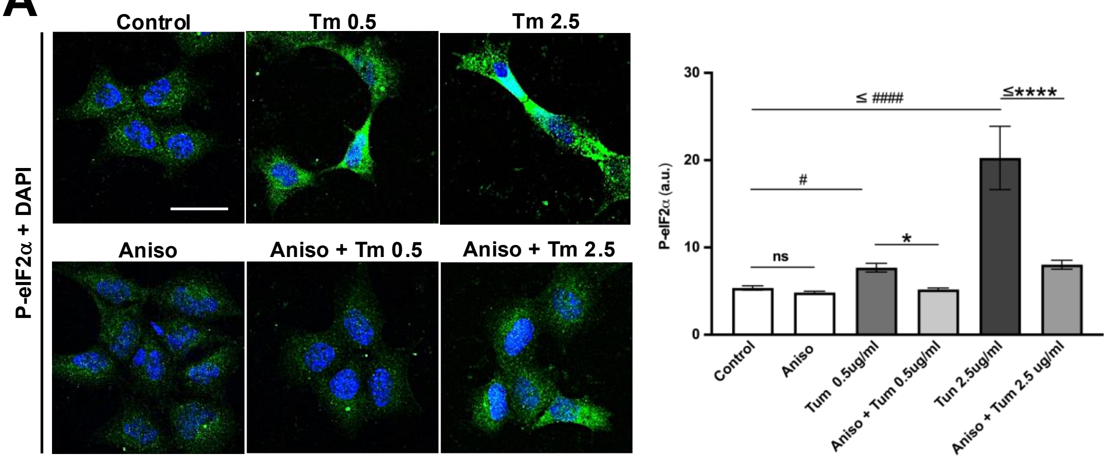

Control
Tm 0.5
Tm 2.5
Aniso
Aniso + Tm 0.5
Aniso + Tm 2.5
P-eIF2α + DAPI
P-eIF2α (a.u.)
30
20
10
0
≤ ####
≤****
#
*
ns
Control
Aniso
Tum 0.5ug/ml
Aniso + Tum 0.5ug/ml
Tun 2.5ug/ml
Aniso + Tum 2.5 ug/ml


B

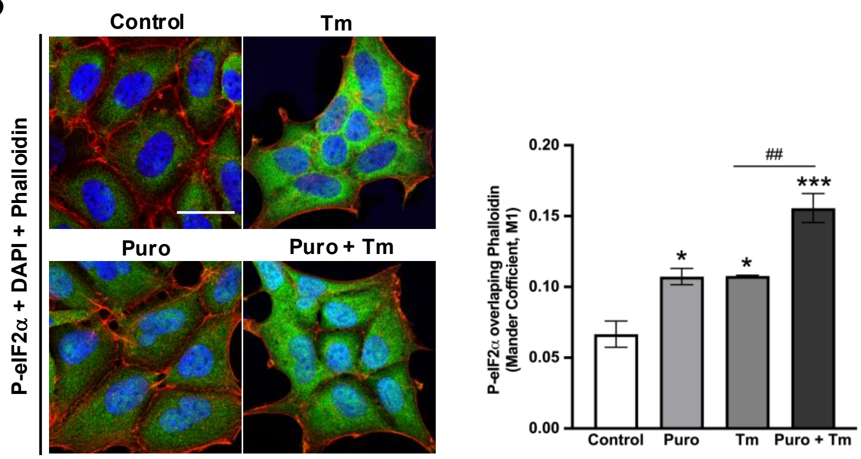

Control
Tm
Puro
Puro + Tm
P-eIF2α + DAPI + Phalloidin
P-eIF2α overlaping Phalloidin (Mander Cofficient, M1)
0.20
0.15
0.10
0.05
0.00
##
***
*
*
Control
Puro
Tm
Puro + Tm


Figure S2

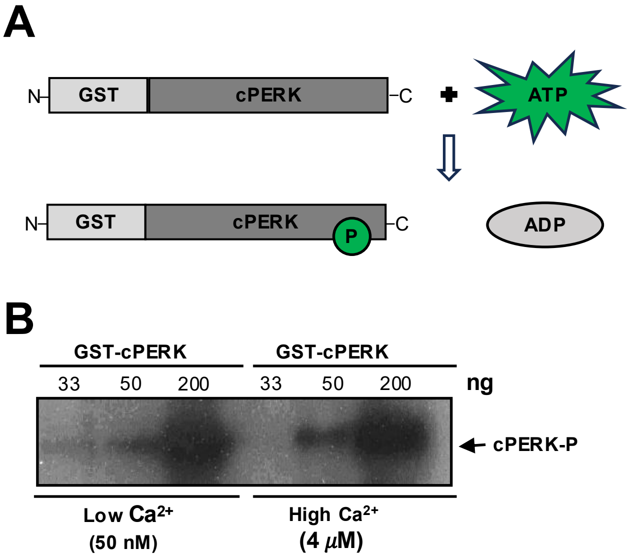
A
N
GST
cPERK
C
ATP
N
GST
cPERK
P
C
ADP
B
GST-cPERK
GST-cPERK
33
50
200
33
50
200
ng
cPERK-P
Low Ca2+
(50 nM)
High Ca2+
(4 μM)

Figure S3

**A**

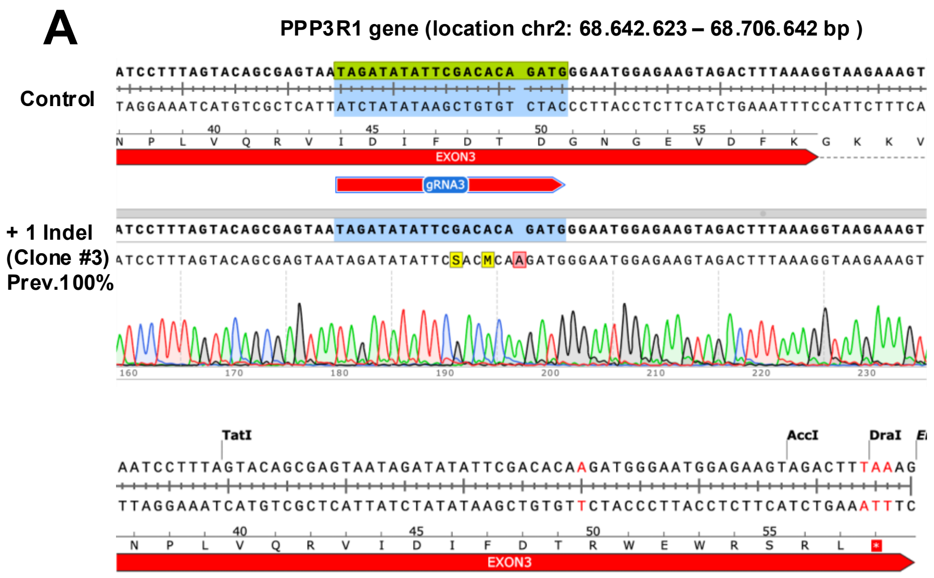


**B**

**CNB**

1 MGNEASYPLE MCSHFDADEI KRLGKRFKKL DLDNSGSLSV EEFMSLPELQ QNPLVQRVID
61 IFDTDGNGEV DFKEFIEGVS QFSVKGDKEQ KLRFAFRIYD MDKDGYISNG ELFQVLKMMV
121 GNNLKDTQLQ QIVDKTIINA DKDGDGRISF EEFCAVVGGL DIHKKMVVDV

**Truncated CNB**

1 MGNEASYPLE.MCSHFDADEI.KRLGKRFKKL.DLDNSGSLSV.EEFMSLPELQ.QNPLVQRVID
61 IFDTRWEWRS.RL*

**C**

**D**

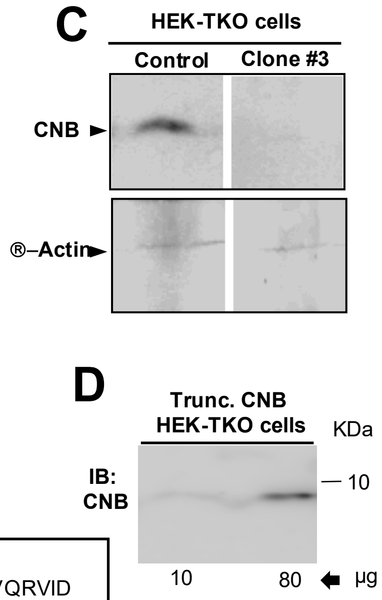


Figure S4

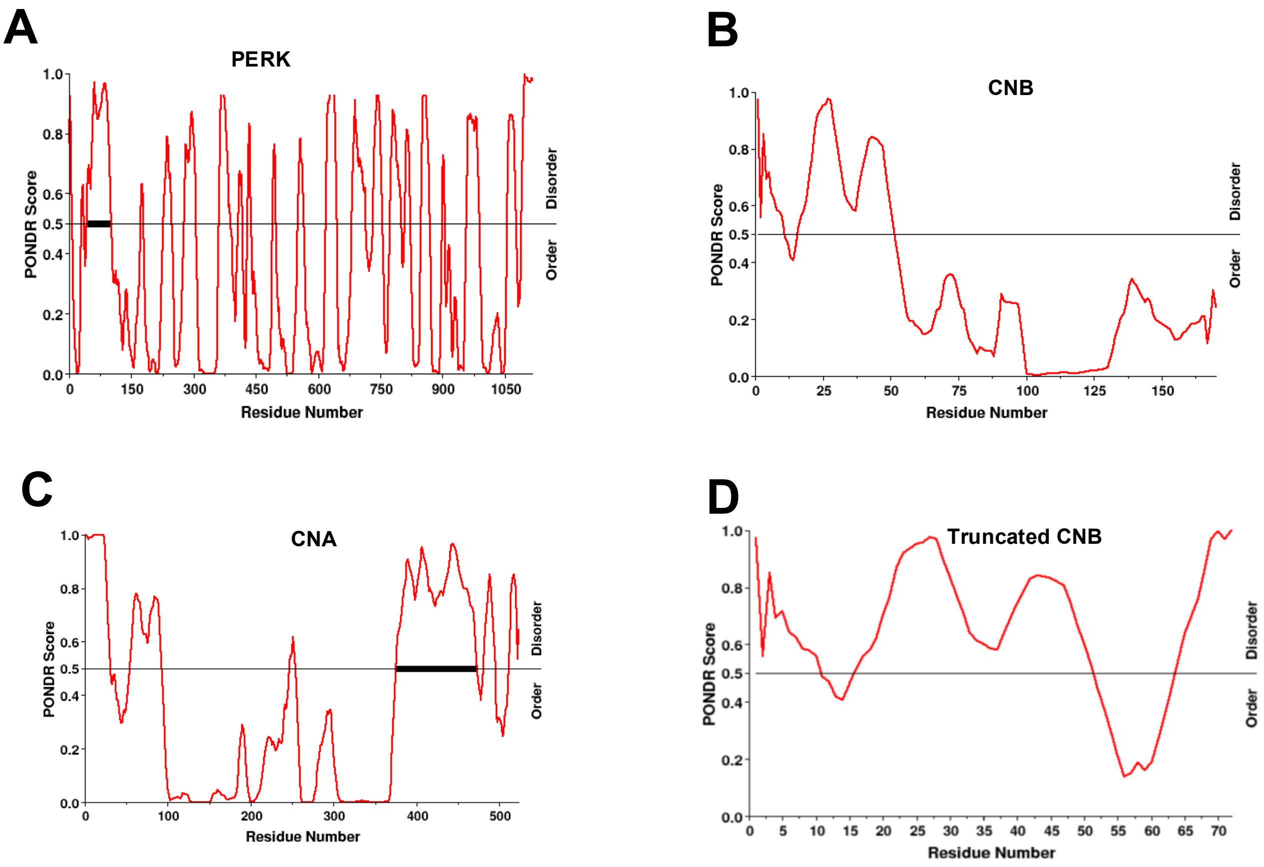
A
PERK
PONDR Score
Residue Number
Disorder
Order
B
CNB
C
CNA
D
Truncated CNB

Figure S5
